# Seasonal influenza viruses show distinct adaptive dynamics during growth in chicken eggs

**DOI:** 10.1101/2025.06.12.659400

**Authors:** Kathryn Kistler, Trevor Bedford

## Abstract

Human influenza viruses are grown in chicken eggs for vaccine production. Sequences of these egg-passaged viruses give us the opportunity to examine the evolution that occurs when these human viruses are subjected to the selective pressure of growing in chicken eggs, which (among other things) express different sialic acid receptors. The repetition of this evolutionary experiment in hundreds of strains over the past several decades allows us to identify mutations that adapt the virus to eggs and epistatic constraints that influence them. We analyze influenza A/H3N2, A/H1N1pdm, B/Vic and B/Yam sequences that were passaged in eggs and find that almost all of the adaptive mutations are located around the receptor-binding pocket of hemagglutinin (HA). We observe epistatic interactions both between adaptive mutations and between these mutations and the continually-evolving human influenza HA background sequence. Our results show that this background dependence is greatest for influenza A/H3N2, then A/H1N1pdm, with B/Vic and B/Yam showing little-to-no background dependence. We find that the total number of adaptive mutations and the length of adaptive walk also follow the same pattern between the influenza subtypes, suggesting that background dependence, number of adaptive mutations, and extent of additive versus epistatic interactions may all be related features of the fitness landscape.

## Introduction

Understanding the fitness landscapes of adaptively evolving pathogens, such as influenza virus, by monitoring their natural evolution is often limited by the fact that we can only observe what happened, and not what could have happened. In other words, by just examining the actual evolutionary trajectory, it is difficult to assess how many possible adaptive pathways were available to the virus and the contingencies of evolution that could influence these pathways. These features of the fitness landscape offer insights about the constraints on evolution.

Here, we analyze what happens when a variety of influenza strains are subjected to the same evolutionary pressure. From this repeated “experiment”, we use strains with different sequences to observe how the background genotype influences possible modes of adaptation and strains with identical sequences to address how many of these different evolutionary pathways are available to a given virus. Specifically, we examine adaptation of human influenza virus to chicken eggs.

The majority of the influenza vaccine doses are grown in chicken eggs ^1^– a practice that was originally developed for military use in response to the devastating 1918 pandemic, and was approved for civilian use in 1945 ^2^. Almost immediately it became clear that the vaccine needed to be reformulated to maintain efficacy against viral antigenic evolution (though the exact cause was not appreciated at the time) and by 1952 the Global Influenza Surveillance and Response System (GISRS) was founded to monitor circulating viruses^3^. As a result of GISRS’s efforts, we have sequences of the 3 seasonal influenza strains that circulate today (A/H3N2, A/H1N1pdm and B/Vic), as well as B/Yam (which recently went extinct in 2020) for the past several decades. These sequences include both direct clinical isolates of the virus and virus that was grown in chicken eggs to produce vaccine.

This temporal collection of sequences tracks the ongoing evolution of influenza viruses in the human population, a process largely driven by antigenic drift. Simultaneously, the egg-passaged sequences reveal the consequences of introducing this changing virus into a different selective environment, namely growth in chicken eggs. In essence, the egg-passaged sequences represent the result of an evolutionary experiment repeated hundreds to thousands of times. During this experiment, we can observe genetic adaptations of the virus as it experiences a relaxation of pressures to maintain human-to-human transmission (including lack of airborne transmission, absence of existing adaptive immunity, and decreased requirement for acid stability to name a few) in addition to a demand for binding different cellular receptors.

Influenza initiates cell entry when its hemagglutinin protein binds to sialic acid moieties decorating cell surface proteins and lipids. Human airways express primarily ɑ2,6-linked sialic acid, while the egg allantoic cavity (where virus is grown) contains ɑ2,3-linked sialic acids. Additionally, human influenza viruses typically bind receptor in the cis conformation, while avian-adapted viruses bind in trans conformation. This difference is known to select for egg-adaptive mutations near hemagglutinin’s receptor binding pocket that increase the virus’ ability to engage ɑ2,3 sialic acid, often at the expense of worse binding to ɑ2,6.

Because the majority of neutralizing antibodies block viral growth by binding to the receptor-binding region of HA (and thus prevent cell entry), mutations that are selected for during egg-passaging can also incidentally affect the antigenicity of the virus. This means not only that the virus grown in eggs can differ from the vaccine strain carefully selected by the WHO, but also that these differences can impact how the virus is recognized by human antibodies ^4^. Low vaccine efficacy has been attributed to egg-adaptation in the 2012-2013 ^5^ and 2016-2017 ^6^ seasons, particularly for the H3N2 component of the vaccine where egg-adaptive mutations have been shown to have the largest impact on immunogenicity ^7^.

Many egg-adaptive substitutions have been described in the receptor-binding domain (RBD) of H3N2 ^8–13^, H1N1pdm ^14–17^, and the human influenza B viruses ^18,19^. Wu et al ^13^ and Liang et al ^20^ have also shown how epistatic interactions in the RBD of H3N2 influence egg-adaptive mutations.

Here, we aim to expand on the evolutionary insights that can be gleaned from the thousands of sequences resulting from growing human influenza isolates in chicken eggs. We take sequences of all available egg-passaged strains and situate them in the context of circulating human virus to determine mutations that occurred during egg-passaging. We do this for all 8 segments of the 4 human influenza viruses (A/H3N2, A/H1N1pdm, B/Vic and B/Yam). We then separate likely adaptive mutations from those found at consensus level due to genetic drift by finding mutations that occurred frequently amongst strains that were independently passaged in eggs. We also consider how different genotypes of human influenza preferentially adapt to egg passaging via different mutations, as well as pairwise epistatic effects between egg-adaptive mutations. We find that the four influenza subtypes differ in their evolution in eggs, and explore what these differences might suggest about the fitness landscapes of these four viruses. Additionally, we assess the number of possible adaptive trajectories available to a given viral genotype, demonstrating the effects of constraint and stochasticity on adaptation.

## Results

### Inference of mutations that occurred during egg-passaging

For each influenza subtype, we downloaded all available egg-passaged sequences dating back to 1995, encompassing 30 years for A/H3N2 and B/Vic, 15 years for A/H1N1pdm (which entered the human population in 2009), and 25 years for B/Yam (which went extinct in 2020). We refer to each human viral isolate, e.g. A/Massachusetts/18/2022, as a strain (meaning it is possible for multiple strains to have the same genotype) and, for this analysis, we use one egg-passaged sequence per strain, where available. This ranged from a minimum of 570 egg-passaged strains for B/Yam to a maximum of 1901 egg-passaged strains for A/H1N1pdm. To determine what mutations occurred during egg-passaging, one could simply compare the sequence of the strain before and after egg-passaging. However, only 37-67% (Figure 1E) of egg-passaged strains were also sequenced prior to egg-passaging. To make use of all of the data, rather than only the egg-passaged strains with available unpassaged sequences, we built phylogenies for each of the 8 segments in the viral genome, positioning egg-passaged strains strains in the context of the circulating human viruses (Figure 1A-D, non-HA segments available at https://nextstrain.org/groups/blab/h3n2/na/egg). This allows us to determine the likely sequence of each strain prior to egg-passaging from the sequence at the internal node ancestral to it in the phylogeny and, thus, to infer mutations that occurred during egg-passaging as those falling on the terminal branch leading to an egg-passaged tip. This allows us to infer egg-passaging mutations for strains that were sequenced prior to egg-passaging (Figure 1F, tip 1) as well as sequences that were not (Figure 1F, tip 3). Because each egg-passaged strain is a separate instance of a human influenza virus being passaged in eggs, we assume that small clades of only egg-passaged strains are actually polytomies.

**Figure 1.**
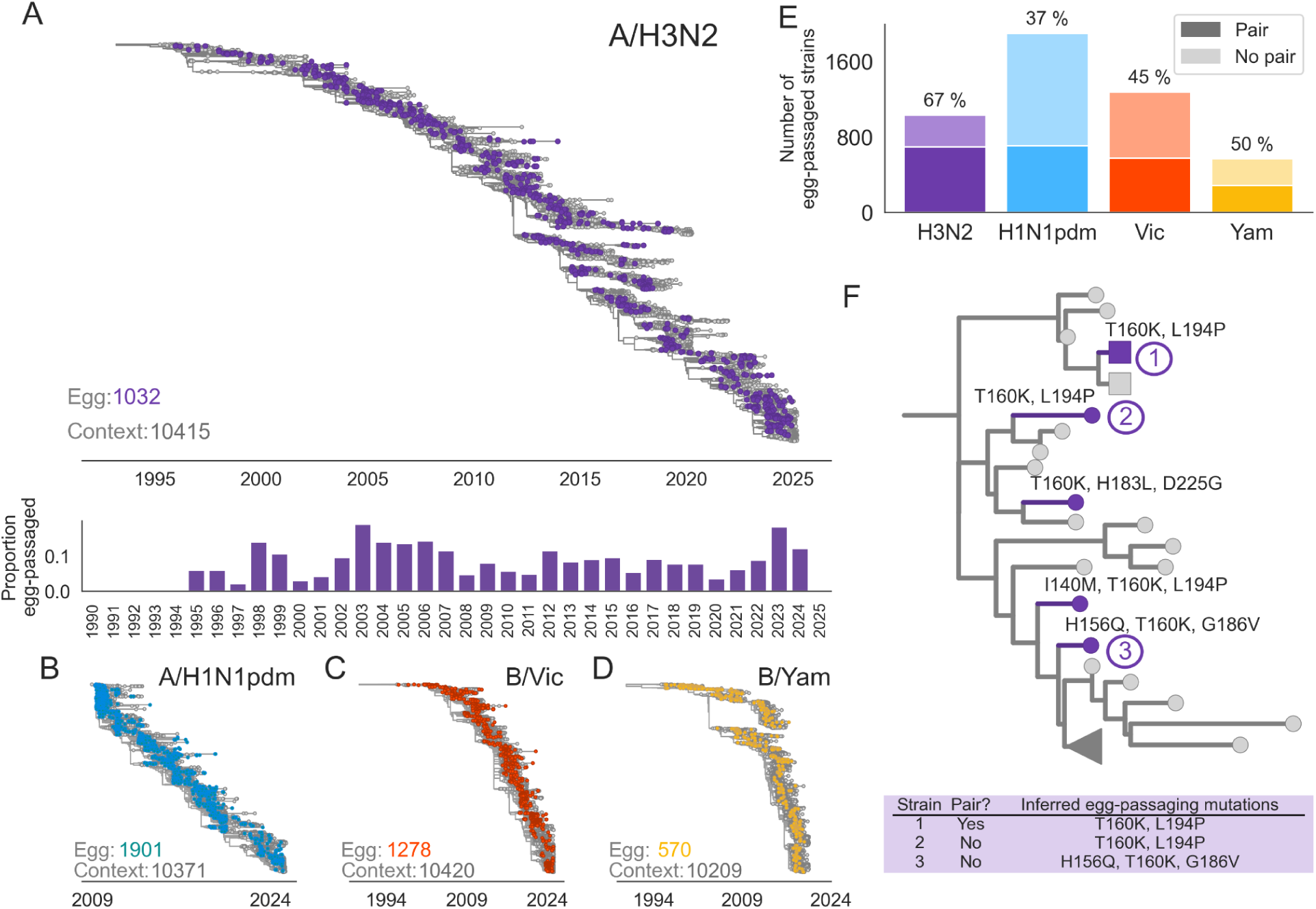
Identification of mutations occurring during egg-passaging of human influenza viruses. A) Phylogeny of 10,415 human influenza A/H3N2 HA sequences (grey), and 1032 egg-passaged strains (purple) sampled between 1995 and 2025. For each year, the proportion of tips that are egg-passaged strains is indicated by the histogram below. B-D) HA phylogenies of egg-passaged strains in the context of non-egg-passaged human influenza virus for A/H1N1pdm (B), B/Vic (C) and B/Yam viruses (D). E) The proportion of egg-passaged strains that were sequenced prior to egg-passaging (pair) is shown for each virus. F) Zoomed view of one clade from the A/H3N2 tree, showing an example of a strain that was sequenced before and after egg-passaging (tip 1), and two strains that were not sequenced prior to egg-passaging where the unpassaged genotype can be inferred from the most closely-related unpassaged strains (tips 2 and 3). The square tip markers indicate a “pair” of sequences of the same strain before and after egg-passaging. Mutations occurring along terminal branches leading to egg-passaged strains are inferred to have occurred during egg-passaging. HA nonsynonymous mutations are labeled along terminal branches.

### Egg-adaptive mutations are primarily located near the receptor binding pocket of HA

Across the eight segments of all four influenza viruses, we observe that 9-40% of strains acquired synonymous mutations during egg-passaging. Longer segments (PB1, PB2, PA) tend to have synonymous mutations in more strains than shorter segments (MP, NS) (Figure 2A), in accordance with expectations of size of target for random mutation. The proportion of strains with nonsynonymous mutations is highest for the HA segment, reaching 81% of strains for H3N2. Additionally, the HA segments of the four subtypes plus the H1N1pdm NP and H3N2 NS segments are the only instances where the proportion of strains with nonsynonymous mutations exceeds the proportion with synonymous mutations.

**Figure 2.**
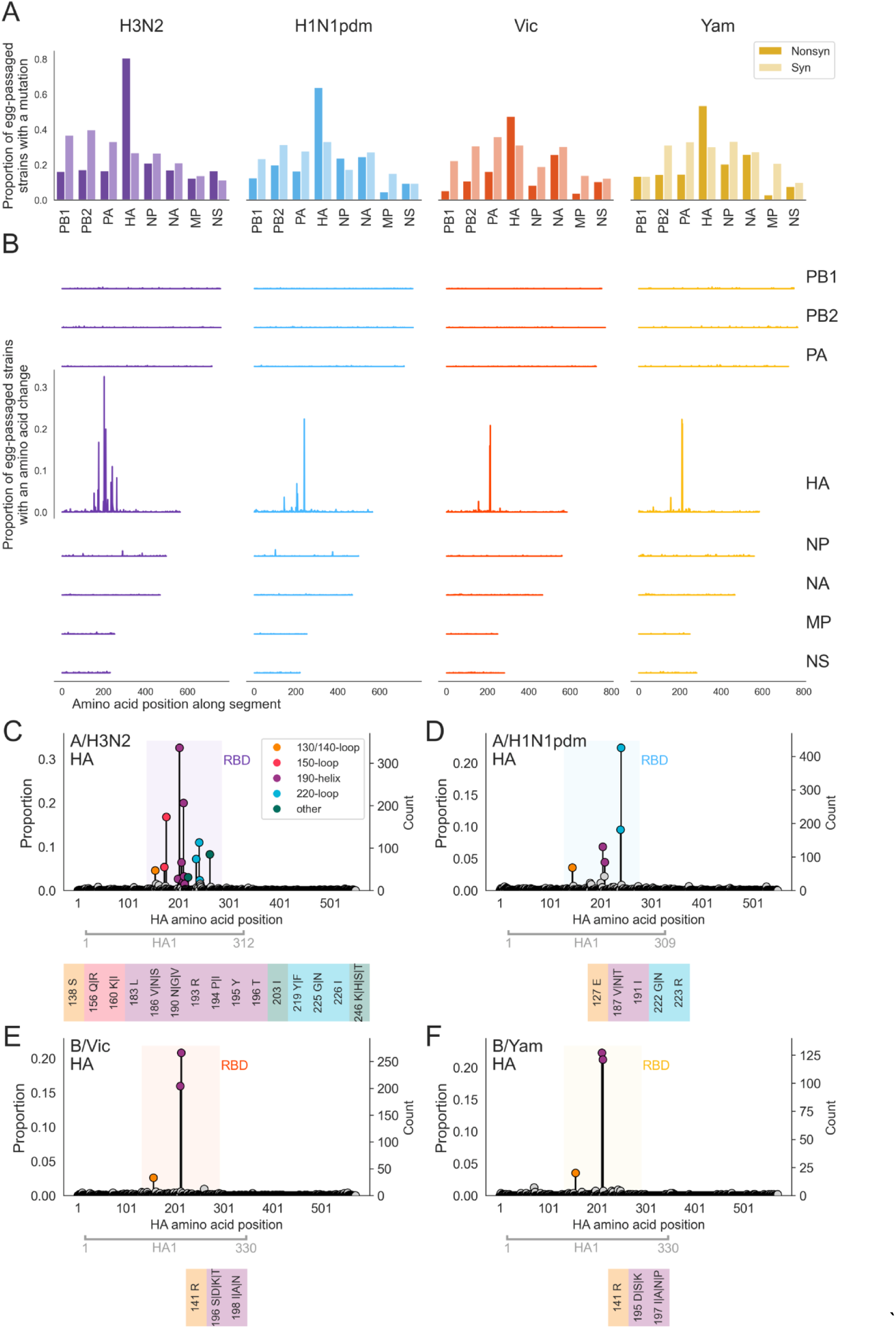
Mutations occur throughout the influenza genome during egg-passaging but amino acid changes are particularly focused in the HA segment, and inferred adaptive mutations are concentrated within the RBD. A) For each influenza subtype, the proportion of egg-passaged strains that got at least one nonsynonymous (dark), or one synonymous (light) mutation during egg-passaging is shown for the coding region of each segment of the influenza virus. B) The proportion of all egg-passaged strains with an amino acid change is plotted at each residue along the coding region of the segment, for each segment. Y-axis and x-axis scales are shared between plots. C-F) The proportion of all egg-passaged viruses that fixed an amino acid substitution is plotted at each residue along the length of the HA protein for C) A/H3N2, D) A/H1N1pdm, E) B/Vic, and F) B/Yam. The location of the RBD is shaded and the positions of inferred egg-adaptive mutations are colored according to whether they fall within a structural feature of the receptor-binding pocket (130 (influenza A) or 140 (influenza B)-loop = yellow, 150-loop = red, 190-helix = purple, 220-loop = blue) or not (green). Below each plot, the residue number and adaptive amino acid substitutions are listed for each adaptive mutation.

We then zoom in to the locations of the nonsynonymous mutations by tallying the proportion of strains with an amino acid change at each residue across the length of the segment. In general, we see an even distribution of amino acid changes across most segments (Figure 2B), indicating that these substitutions are occurring at random, rather than at functionally-important residues. In stark contrast to the flat topography of these plots for most segments, we see tall spikes within the HA histograms, marking residues that were independently mutated in many strains, and suggesting that amino acid changes at these positions have been selected for.

Each egg-passaged strain in this analysis is the result of passaging a different strain in chicken eggs, meaning that we have not included multiple sequences of serial passages of the same strain. Therefore, each sequence is an independent recounting of the evolutionary consequence of growing human influenza virus in chicken eggs. Because of this, the number of times a particular mutation is observed in different strains is reflective of its selective coefficient. In other words, a mutation occurring just once across hundreds of egg-passaged strains is likely evolutionarily neutral in this context, while a mutation that appears in many strains has likely been selected for due to a fitness advantage in the face of pressure common to all of these strains, i.e. growth in chicken eggs.

Using this logic, we separate egg-adaptive mutations from neutral and deleterious mutations by the number of times they occur amongst all egg-passaged strains. Specifically, we pick out residues that were mutated in at least 2.5% of egg-passaged strains overall, or specific substitutions that reach at least 7.5% within a 3-year window. We believe this threshold is fairly conservative, allowing us to be confident that the mutations we have identified are truly adaptive, but at the cost of likely missing some mutations with only a small beneficial effect. By this metric, all putatively egg-adaptive mutations occur in HA or NP.

We identify residues with egg-adaptive mutations within the HA segment of all four viruses (Figure 2C-F). Every one of these sites is located within the receptor-binding domain of HA1, and most of them are found within structural features of the HA protein that surround the receptor-binding pocket (130-loop, 150-loop, 190-helix and 220-loop). The influenza B adaptive mutations are primarily focused in the 190-helix, while the influenza A (especially H3N2) mutations are more spread out between these features. In fact, all of the Vic adaptive mutations occur at structurally homologous positions to the Yam mutations, and the amino acid substitutions observed amongst adaptive mutations are also almost identical (Supplemental Figure 1). The influenza A viruses share two structurally homologous loci of egg-adaptation in the 190-helix and another two in the 220-loop, again, getting mostly the same amino acid substitutions. The four egg-adaptive residues shared between H3N2 and H1N1pdm were also identified by Dadonaite et al ^21^ as being crucial for ɑ2,3-versus ɑ2,6-sialic acid binding in H5 viruses.

Highlighting the egg-adaptive sites on the structure of HA emphasizes the proximity of these residues to the receptor-binding pocket (Supplemental Figure 2). The striking enrichment of mutated sites flanking the receptor-binding pocket of these viruses suggests that there is strong selection for protein change at the interface of hemagglutinin and the sialic acid receptor, likely to improve binding of the altered conformation of these sugars presented by avian egg cells compared to human respiratory cells. Some of these mutations are known to directly impact binding of human versus avian sialic acid ^13,14,22^, while others have been noted to abolish glycosylation ^6,23–25^. Egg-adaptive mutations that prevent glycosylation include the 196/198 (or 195/197) mutations in Vic (or Yam), and the H3N2 mutations at 160 and 246. Though we cannot determine the reason mutations that remove glycans from HA1 are selected for in eggs, it seems likely that either these glycans compromise binding to ɑ2,3-sialic acid and/or a pressure retain glycans as a defensive shield against human neutralizing antibodies is relaxed.

We also detect a few potentially adaptive mutations in the nucleoprotein (NP) segment of both influenza A viruses (Supplemental Figure 3). H3N2 viruses have two residues in NP that are repeatedly mutated during egg-passaging (290 and 384), while H1N1pdm strains have three sites (101,102 and 375), including two neighboring residues. Fitting with a potential role in host adaptation, all 5 of these residues are found amongst the sites that Kukol and Hughes identify as varying most between hosts ^26^. The NP protein is essential for transporting viral RNA into the nucleus for replication ^27^, a process that relies on interactions with importins and other host cell machinery that differ between humans and birds ^28^. However, the positions where we observe mutations do not fall within NP’s nuclear localization sequences ^29^, indicating that egg-adaptation may be through a different mechanism. Chen et al ^30^ also found that nucleoprotein has the most positions out of any segment that act as signatures for differentiating human from avian influenza strains, but the mutations we observe are also not amongst these sites.

### The circulating human influenza genotype influences which egg-adaptive mutations can occur

Human influenza viruses have undergone a considerable amount of antigenic evolution during the past 30 years and, as with egg-adaptation, antigenic evolution is also focused in the receptor-binding domain of HA1. Thus we expect that egg-adaptive mutations will change over time as the human virus evolves, either due to the human virus incidentally becoming more or less fit in eggs or because of epistatic constraints of the HA structure. Plotting the prevalence of mutations at each adaptive residue over time reveals that some mutations to some sites are temporally restricted (e.g. H3N2 225 and H1N1pdm 191), while others are not (e.g. H1N1pdm 223 and Yam 195) (Figure 3, left). Additionally, at sites where we observe adaptive changes to multiple different amino acids, we again see that some substitutions, such as H3N2 194I vs. 194P, are time-dependent while others, like H3N2 219F vs. 219Y, are not (Figure 3, right).

**Figure 3.**
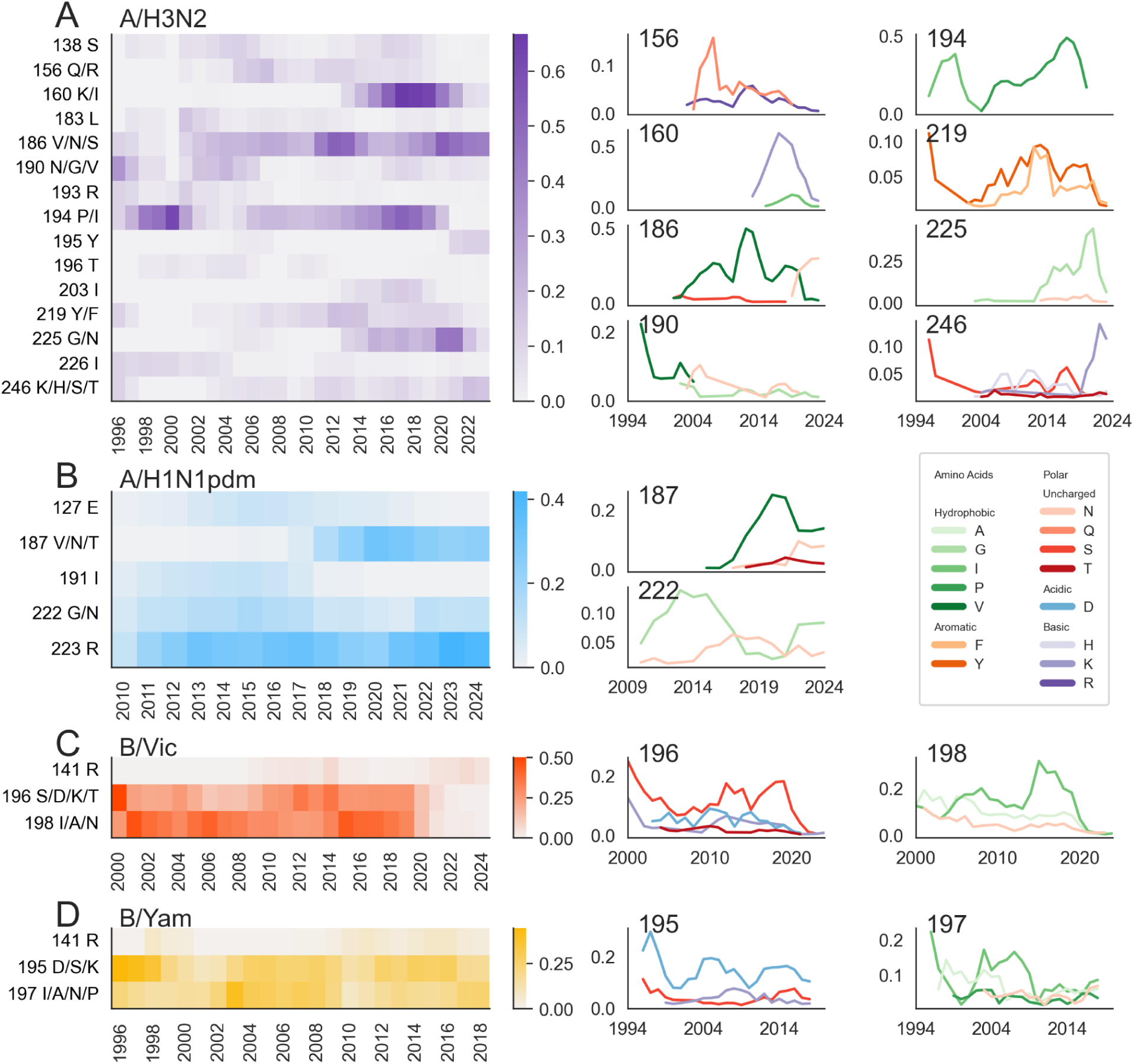
Egg-adaptive mutations change over time as the human influenza virus evolves. The proportion of egg-passaged strains with a mutation at each egg-adaptive residue is calculated in 3-year sliding windows and plotted as a heatmap for A) H3N2, B) H1N1pdm, C) Vic, and D) Yam. For each residue where substitutions to multiple amino acids are observed, the frequency of each of those substitutions is plotted over time.

#Figure 3 reveals differences in the time dependence of egg-adaptive mutations between the different influenza subtypes. Though some mutations are observed for a decade or longer while others are confined to just a handful of years, we observe temporal restriction for every single H3N2 egg-adaptive site, and among many of the specific adaptive substitutions at these sites as well. The other influenza A virus, H1N1pdm, has some egg-adaptive mutations that are prominent across the entire time frame (223R and 222G/N) while the mutations at the other three sites occur only during a section of time. It should be noted that this analysis spans 30 years for H3N2 and only 15 years for H1N1pdm, and it is possible that H1N1pdm 222 and 223 mutations will not continue to be egg-adaptive on a longer time scale. In contrast to the influenza A viruses, influenza B/Yam shows no time dependence, and the dominant B/Vic sites (196 and 198) get mutated regularly and consistently until around 2020 when mutations to both sites stop appearing.

Time dependence of egg-adaptive mutations is a proxy for dependence on the genotype of the human influenza virus that was passaged in eggs. To determine which genetic changes that occurred during the evolution of these influenza viruses as they circulated in humans subsequently impacted egg-adaptation, we identified HA1 substitutions occurring on the trunk of the tree that correspond best with presence or absence of a particular egg-adaptive mutation. We are able to identify genetic changes in the human virus that correlate with many of the time-restricted egg-adaptive mutations. Based on these results, we classify the genetic dependence of egg-adaptive mutations on human genotype into three main categories that likely explain the time-dependence: 1) fixations in the human virus that alter fitness in eggs, 2) mutational codon accessibility, and 3) epistatic interactions between the adaptive mutations and the background. The first category includes mutations like H3N2 186 and Vic 196 mutations (Figure 4), where the human virus fixes substitutions at the egg-adaptive site, resulting in multiple genotypes with potentially different fitnesses in eggs.

**Figure 4.**
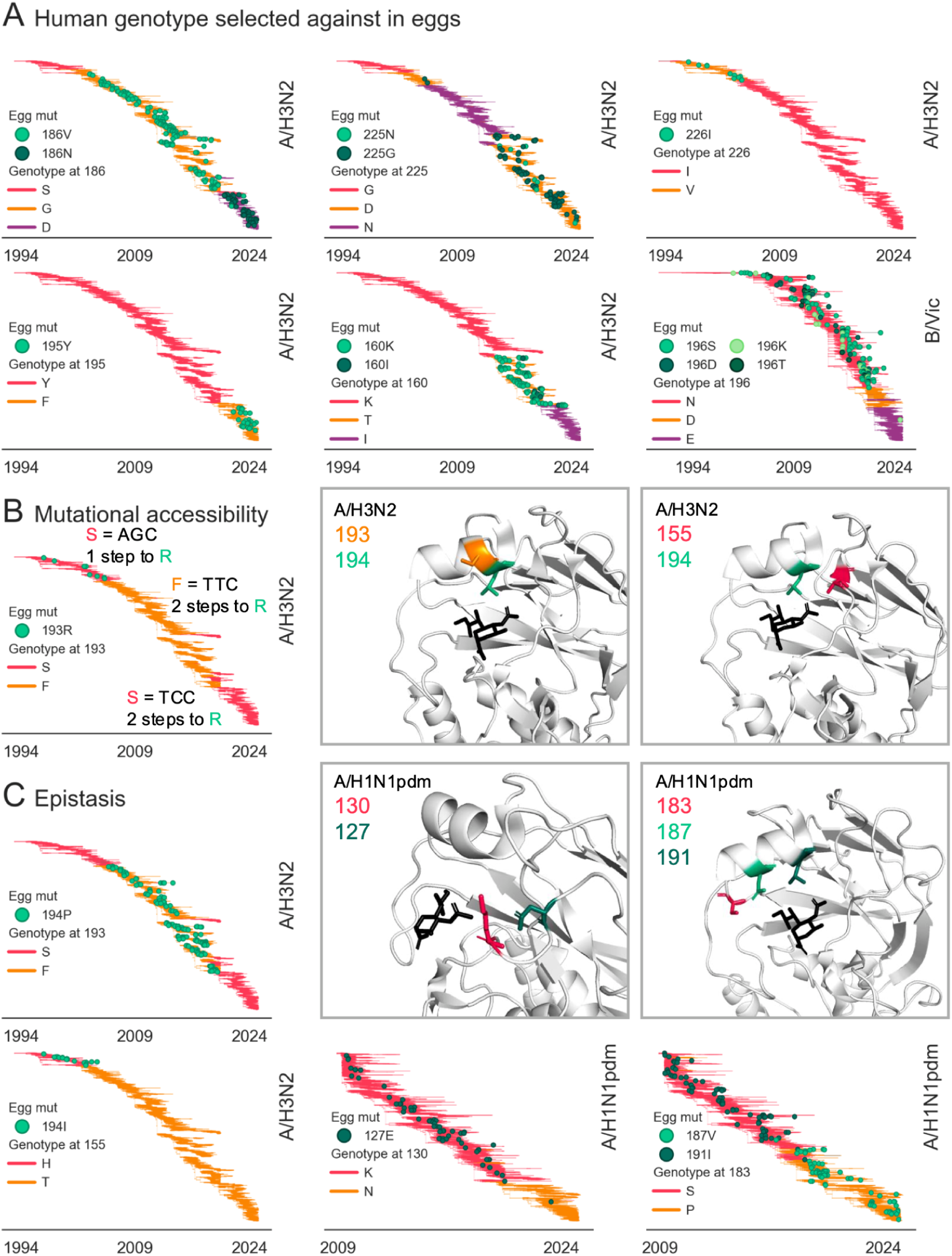
Egg-adaptive mutations depend on the genotype of the human influenza virus. The distribution of egg-passaged strains with adaptive mutations (green circles) is shown on top of the genotype of the circulating human influenza virus (colored branches). Influenza subtype is labeled on the right of each plot. Temporal restriction of adaptive mutations can depend on the human virus’ genotype at the position of egg-adaptation via selection against the human genotype (A) or mutational accessibility (B), or it can depend on the genotype at other positions in HA1 via epistasis (C). The location of epistatically-interacting residues is highlighted on the structure of HA in the insets.

As seen in Figure 4A, human H3N2 had a serine at position 186 until around 2000, when 186G swept. We observed no egg-adaptive mutations on top of 186S (encoded by codon AGT), even though the highly-adaptive 186N is accessible via a single nucleotide mutation, indicating that 186S has high fitness in eggs. Once 186G sweeps, we begin to see frequent mutations to 186V during egg-passaging, with as many as 63% of egg-passaged strains during a single year (2013) getting 186V (Figure 3A). During this time period, there are also a handful of egg-passaged strains that get 186S, again pointing towards 186S being fairly high fitness in eggs. The vast discrepancy between the number of egg-passaged strains that get G186V vs. G186S speaks to valine being even more beneficial for egg growth than serine during this time period. The 186N mutation was not available during this time period, but following the subsequent fixation of 186D in the human virus, both egg-adaptive mutations (186N and 186V) become accessible via a single nucleotide mutation. During this time period, 38% of the 245 egg-passaged strains get the 186N mutation while 186V occurs in just 5 strains. In the absence of epistatic influences from other sites throughout RBD, this suggests that H3N2 has a preference for N, then V, then S at 186 when grown at eggs, and that G and D have fairly low fitness, though epistasis likely does complicate this conclusion.

We can similarly work out the relative fitnesses of different genotypes at other HA1 positions based on the genotype of the human virus and the occurrence and identity of observed egg-adaptive mutations. For instance, the circulating human Vic virus has 196N (encoded by the AAC codon) from 1994 until around 2020, and every observed egg-adaptive mutation at 196 occurs during this period. Around 2020, 196D fixes in the human population and egg-adaptive mutations at this site cease, which makes sense because 196D is actually one of the observed egg-adaptive mutations, further suggesting that 196N has lower fitness in eggs than 196D. After a matter of just a couple years, the circulating virus again fixes a 196 mutation, changing the human virus to 196E (encoded by GAA). Though 196E is not an egg-adaptive mutation we observed, we suggest that it is also high fitness in eggs because we observe only a single egg-adaptive mutation on human strains with 196E, despite the fact that both D and K adaptive substitutions are mutationally accessible. We suggest that the reason E is not observed after egg-passaging during the long span of time when egg-adaptive mutations are occurring on 196N is because E requires two mutations from AAC (encoding N), whereas the other adaptive mutations are accessed by just one.

With H3N2 193R, we see a strong example of how distances between codons can impact whether or not an adaptive mutation will occur. Until around 2004, circulating human H3N2 had a serine encoded by AGC at residue 193, and during this period we observe several egg-passaged strains mutating to 193R during this period. However, in the ensuing years, a couple nucleotide mutations fix in the 193 codon, resulting in a sweep of phenylalanine and then a reversion to serine. The serine reversion is encoded by TCC, which is now two mutational steps away from R, emphasizing that despite evidence that 193R is preferred over 193S in eggs, it will not occur due to mutationally accessibility. The reason accessibility matters so much is because a substitution requiring two mutations would either need to get both precise mutations simultaneously (which is very unlikely) or acquire a first mutation followed by the second (which is also very unlikely since the first mutation would not be selected for).

Finally, we also see that some egg-adaptive mutations are impacted by epistatic interactions with other sites in the HA protein (Figure 4C). An example of this is H3N2’s egg-adaptive 194P mutation, whose occurrence is bounded tightly by a phenylalanine at position 193 in the human virus. We see 194P occurring exclusively on top of 193F, and never on 193S. This includes a couple of small clades with 193S in the 2010’s that never developed 194P mutations when passaged in eggs, while their sister clades with 193F did.

The H1N1pdm 191I mutation occurs in eggs from 2009 until 2017-2018, around which point egg-adaptive mutations at residue 187 started occurring (Figure 3B). This switch coincides with a S-to-P fixation at position 183 in the circulating human virus. Of the 86 egg-passaged strains with a 191I mutation, all but two occur in human strains with 183S, and of the 125 egg-passaged strains that acquired 187V/N/T mutations, 100% occur on top of 183P (Figure 4C). Both 187 and 191 are located on the side of the 190-helix facing the receptor-binding pocket whereas 183 is located in the loop preceding this helix. Because prolines limit rotation around the protein background, we hypothesize that the S183P fixation may alter the position of the 190-helix such that mutations at residue 191 are no longer able to improve receptor-binding to ɑ2,3-linked sialic acids, and instead, mutations at 187 become important for this function. In support of this theory, the equivalent residue H3N2 (186) is a major site of egg-adaptation and a mutation from G to V at this site was shown to shift the location of the 190-helix, thereby expanding the receptor-binding pocket ^13^.

### Epistasis between egg-adaptive mutations influences the length of adaptive trajectories

We next consider how the mutations acquired during influenza’s pursuit of egg-adaptation are influenced not just by the genotype of the human virus injected into the egg, but also by the presence of other egg-adaptive mutations. For each pair of egg-adaptive mutations in HA1, we calculate an enrichment ratio to determine whether the mutations co-occur more or less often than would be expected from the individual frequencies of the two mutations. To account for background dependency, we require that the period of time the two mutations are observed must overlap by at least three years. As seen in Figure 5A, the H3N2 plot has both many grey boxes-recapitulating strong background dependency of H3N2 mutations shown by Figure 3A- and many more red boxes, indicating that egg-adaptive mutations co-occur more commonly in H3N2 than they do in H1N1pdm or influenza B viruses. In fact, egg-adaptive mutations never co-occur in influenza B viruses (Figure 5C-D). In egg-passaged H1N1pdm viruses, there is also a dearth of co-occurring adaptive mutations, with the neutral enrichment scores primarily being observed between mutations at 222 and another residue (Figure 5B).

**Figure 5.**
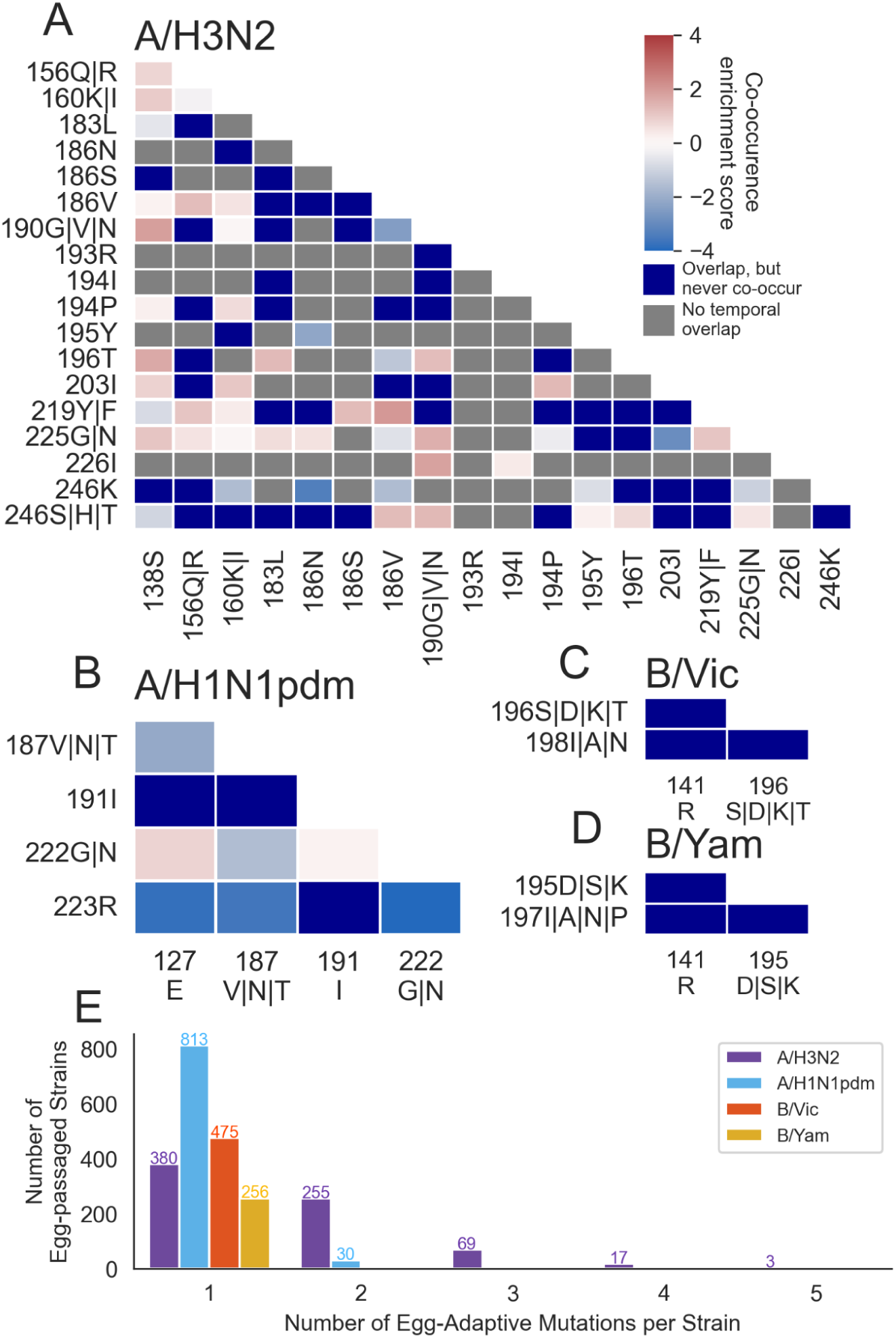
Epistatic interactions between egg-adaptive mutations influence their co-occurrence. A) A mutational co-occurrence enrichment score is calculated for each pair of mutations that overlaps temporally as the ratio of observed to expected mutational co-occurrence, specifically by 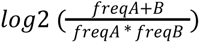 where A and B are the two mutations. Mutations that co-occur more often than expected based on their individual frequencies will have higher (redder) values, and mutations that co-occur less frequently have lower (bluer) values. Mutations that overlap temporally but never co-occur together are shown in dark blue because *log*2 of 0 is undefined. B) The total number of egg-adaptive HA mutations per strain is plotted by influenza subtype. Only strains that get at least one adaptive mutation during egg-passaging are plotted.

The mutually exclusive nature of egg-adaptive mutations in Vic and Yam indicates that as human influenza B virus adapts to eggs, it has access to at least 3 distinct fitness peaks, separated by sign epistasis. The source of this epistasis could either be because the mutations are structurally incompatible or because they have completely non-additive beneficial effects. In the latter case, it would be possible that the mutations have completely overlapping effects or that their mechanisms of adaptation are actually antagonistic to each other. Both of the 190 mutations act to remove the same glycan, so the presence of one of these adaptive mutations drops the effect of the other to neutral, or even deleterious. It is difficult to tell from these data why 141R does not co-occur with either 190-helix mutation.

In contrast, the pairwise co-occurrence of some H3N2 mutations with each other suggests that egg-adaptation in this virus can be achieved through the additive, or even synergistic, effects of multiple mutations. In particular, we see that some adaptive mutations like 138S, 160K/I and 225G/N co-occur with other mutations frequently while others, like 246K and 186N do not. The fixation of multiple adaptive mutations in the same strain suggests either that the mutations have distinct, non-overlapping beneficial effects for the virus, or that mutations actually have similar molecular benefits that can act in coordination. Others have provided evidence that both of these are at play in H3N2 egg-adaptation. The mutations at residue 160 remove a glycan and it is likely that the advantage of this is independent of mutations that alter the receptor-binding pocket itself. On the other hand, the combination of 186V and 226I increased viral growth in the egg allantois where neither mutation alone did, implying a synergistic effect of these mutations ^9^.

However, negative epistasis also creates distinct fitness peaks during H3N2 egg-adaptation. Figure 5A shows a depletion of 186N co-occurring with 195Y mutations. Circulating human H3N2 had a tyrosine at residue 195 until phenylalanine fixed around 2020. After this point, 186N mutations and reversions to 195Y both become frequent in egg-passaged strains, occuring in 94/275 and 29/275, respectively. However, these mutations only occur together in the same strain twice, indicating that 186N and 195Y both strongly increase fitness in eggs but that their benefits are not additive. Similarly, 186V and 194P mutations are both very common from 2005-2019, but are never found simultaneously. Wu et al ^13^ have shown that mutual exclusivity of these highly adaptive mutations is due to a structural incompatibility. These results suggest that, like for influenza B viruses, the fitness landscape of egg adaptation by human H3N2 also has multiple peaks separated by sign epistasis but, unlike influenza B viruses, the H3N2 peaks contain longer adaptive walks consisting of multiple adaptive mutations, rather than just one.

This trend toward longer adaptive trajectories (or more mutational co-occurrence) in H3N2 is again evident when assessing the total number of adaptive mutations per egg-passaged strain (Figure 5E). Of H3N2 strains that get adaptive mutations in HA1 during egg-passaging, 48% of them get multiple, with some strains getting 3, 4 or even 5 adaptive mutations. This is in stark contrast to 4% of H1N1pdm strains that get 2 mutations and 0% of Vic or Yam getting multiple egg-adaptive mutations.

If we view H3N2 and the influenza B viruses as having different modes of egg-adaptation with regards to epistasis between adaptive mutations and the length of adaptive trajectories, H1N1pdm is somewhere in between.

### Egg-adaptation of human influenza virus reveals many possible adaptive trajectories

The sign epistasis we observe in Figure 5 suggests that a given influenza hemagglutinin protein can access multiple available pathways towards egg-adaptation. Figures 3 and 4 demonstrate that the pathways available to a given hemagglutinin change over time due to background dependence for H3N2 and to some extent H1N1pdm but very little for influenza B strains. This suggests that, amongst egg-passaged strains derived from identical unpassaged strains, we will not observe exact repetition of egg-adaptation, but rather an assortment of the different adaptive mutations available to that background. To test this more directly, we compare egg adaptation between sets of strains that have identical HA1 sequences before egg-passaging. We identify 37 sets where at least 5 egg-passaged H3N2 strains derive from the same unpassaged HA1 sequence (48 H1N1pdm sets, 34 Vic sets, and 23 Yam sets).

Indeed, we see that when a virus with a given HA1 sequence is passaged in eggs many times in parallel, the virus will not adapt to eggs via the same adaptive pathway each time. For H1N1pdm and influenza B viruses, we see many examples where the same HA1 sequence will develop each of the possible egg-adaptive mutations in independent replicates (Figure 6A, Supplemental Figures 5-7). Again, we observe only single adaptive mutations occurring in influenza B viruses and just a handful of H1N1pdm viruses with two egg-adaptive mutations. For H1N1pdm, we observe that HA1 sequences that have access to an adaptive mutation at residue 187 do not have access to 191 and vice versa. In general, the number of different adaptive mutations observed within a set increases with the number of strains in that set (Figure 6B). This supports our hypothesis that each of these influenza viruses have access to multiple mutually exclusive egg-adaptive pathways, and which mutation fixes will depend on the relative selective coefficient of that mutation and, importantly, to stochasticity of the mutation occurring.

**Figure 6.**
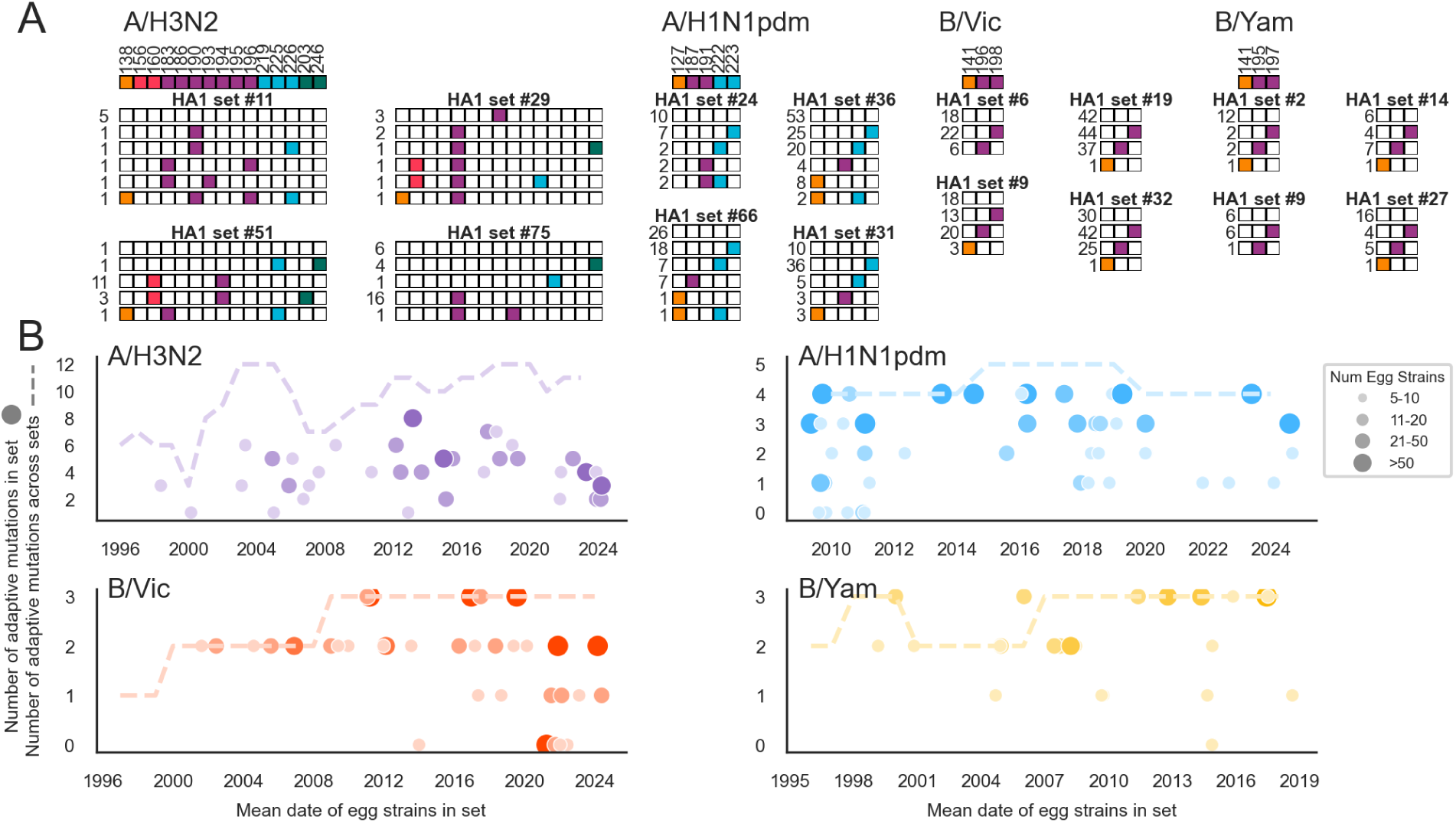
A given hemagglutinin sequence has access to multiple possible egg-adaptive trajectories. A) Four representative sets of egg-passaged sequences derived from identical unpassaged HA1 sequences are shown for each virus. The “adaptive haplotypes” observed within a set are represented by a string of boxes that are filled in if there was an adaptive mutation at that position. For each subtype, a key denoting which column of boxes corresponds to which position is shown at the top. The number of egg-passaged strains in the set with that “adaptive haplotype” is written to the left. All other sets can be seen in Supplement Figures 4-7. B) The number of adaptive sites mutated amongst each set is plotted over time, with the circle size indicating how many egg-passaged strains are in the set. Each set is plotted at the mean date of all strains within that set. The dashed line indicates the total number of adaptive positions observed throughout all egg-passaged sequences over time (see Supplemental Figure 9).

The H3N2 sets reflect a higher degree of background dependence as noted in Figures 3 and 4, with only a subset of adaptive mutations appearing amongst the egg-passaged strains in each set. There are several H3N2 sets where many viruses fixed the same mutation, but some viruses also got an additional mutation or two, suggesting incomplete evolution down a shared adaptive pathway. Perhaps if the viruses were grown in eggs for longer, the results of egg-adapted H3N2 viruses would converge even more. Potentially incomplete adaptation, strong background dependence, and H3N2’s faster rate of evolution in humans (resulting in smaller set sizes, on average) could all contribute to the observation that we never see H3N2 sets with mutations at every site that is observed amongst all egg-passaged strains during the same time period. This is shown by Figure 6B, where in contrast to H1N1pdm, Vic and Yam, all H3N2 sets fall below the dashed line.

Being able to look at repetition of the same hemagglutinin sequence adapting to chicken eggs over and over again gives us a unique view into what the possibilities of evolution are. In other words, we can infer that an H1N1pdm strain that adapted to eggs via a 191I mutation could have instead adapted via a mutation at 127, 222, or 223, and the fact that 191I is what occurred is due, in large part, to stochastic effects. With this framing, we can consider the differences between avian influenza A and human influenza A viruses. We hypothesize that egg-adaptation should mirror differences between human and avian HA that enable them to better bind ɑ-2,6 versus ɑ-2,3-sialic acids.

For H1 and H3, we compare the genotypes observed in human and avian influenza at each egg-adaptive site. We notice that, at most sites, the egg-adaptive amino acid is not what is observed in avian influenza, and in fact, that avian and human HAs often have the same or similar amino acids at these sites (Supplemental Figure 8). However, avian H1 has 127E and the highly egg-adaptive 222G and avian H3 has genotypes 186S, 195Y, 225G that we observe to be egg-adaptive. We speculate that these sites represent the different pathways avian and human influenza hemagglutinin evolution have actually taken to recognize ɑ-2,3 versus ɑ-2,6-sialic acid, while the results from our egg-adaptation analysis represent alternative evolution that could have taken place to differentially allow these hemagglutinins to better bind the sialic acids of their host species.

Notably, the human H3N2 virus has fixed each of these egg-adaptive mutations that are found in avian H3 (186S, 195Y, and 225G) at different points during the past 30 years (Supplemental Figure 8). This is not the case for H1N1pdm: we don’t observe egg-adaptive mutations fixing in the human virus population. This suggests that during its ongoing evolution in humans, the human H3N2 virus incidentally becomes more and less able to bind avian ɑ-2,3 sialic acid, a hypothesis supported by a fluctuating total number of adaptive mutations over time (Supplemental Figure 9).

## Discussion

In the study of viral evolution, say that of influenza viruses, we know stochasticity and historical contingency are influential forces, yet we are often limited to scrutinizing what did happen, and it is much harder to assess what could have happened. This leaves open important questions regarding how determinative versus stochastic adaptive evolution is; does the virus have 100s of possible adaptive trajectories? Tens? A couple? One? Natural populations of human viruses are under a wide and changing range of selective pressures, making it quite complicated to determine all of the possible evolutionary trajectories. Here, we looked at a less complicated and more tightly bounded selective regime: adaptation of human influenza virus to chicken eggs. By assessing repeated evolution in this situation, we found that human influenza virus has a handful of different adaptive trajectories toward egg-adaptation, that these trajectories change as the human virus evolves, and that all of this differs between the four human influenza subtypes.

We find that, during egg-adaptation, influenza subtype A/H3N2 exhibits the most background dependence, the most positively-epistatic interactions, and the longest adaptive walks. In all of these aspects, A/H1N1pdm ranks second, followed by the two influenza B subtypes, which adapt very similarly. These findings are consistent with at least two different (non-mutually exclusive) hypotheses to explain the differences in egg-adaptive evolution between the subtypes: 1) human influenza B viruses are more fit in eggs, and/or 2) human influenza A viruses (and particularly H3N2) have more adaptable hemagglutinin proteins. In support of hypothesis 1, more possible adaptive mutations and longer adaptive walks (as seen in H3N2) are characteristics of a population further from a fitness peak, while mutually exclusive adaptive mutations mediated by sign epistasis (like Vic and Yam) are characteristic of a population close to a fitness peak ^31–33^. This hypothesis seems quite plausible except that influenza A viruses (including H3NX and H1NX) are prolific in wild bird populations, while birds are not a natural host of influenza B viruses. On the other hand, some proteins are known to be more generally tolerant to mutations than others, and it is possible that the influenza A hemagglutinins are more flexible in a way that allows them to more easily fine-tune the receptor-binding pocket to accommodate different sialic acids. Interestingly, this same ordering of subtypes also describes their relative rates of antigenic drift ^34,35^, offering the possible hypothesis that this H3 hemagglutinin has more mutational flexibility in the receptor-binding pocket, which allows it to more readily adapt to ɑ2,3 receptors and also to more readily access antigenic variation that does not compromise receptor-binding. From our results, we cannot distinguish between the potential explanations of the different egg-adaptation observed between influenza subtypes.

We also find that the total number of egg-adaptive sites varies more over time for A/H3N2 than it does for the other viruses (Supplemental Figure 9). This result aligns with H3N2’s higher background dependence (Figures 3 and 4), and suggests that evolution of human H3N2 brings the virus into different regions of the fitness landscape in eggs. The ongoing antigenic evolution in humans incidentally makes the virus more or less fit when it is subsequently passaged in chicken eggs, and also can alter available pathways towards egg-adaptation (Figures 3-6).

In the context of a rapidly-evolving protein that fixes ∼0.5-3 amino acid substitutions in HA1 subunit each year, we find the degree of parallelism between egg-adaptation of the different influenza subtypes to be quite striking. The influenza B lineages, Vic and Yam, diverged in 1983 ^36^ and since then, their hemagglutinin proteins have been undergoing separate antigenic evolution including fixations at different residues and a deletion in B/Vic ^37,38^. Despite this, we observe that both viruses adapt to eggs via similar substitutions at equivalent residues (Supplemental Figure 1). The human influenza A subtypes are estimated to have diverged hundreds or thousands of years ago ^39^, yet they share several prominent sites of egg-adaptation (Supplemental Figure 1), and these sites were also shown to contribute to ɑ2,3 versus ɑ2,6 receptor preference in H5 ^21^ and H7 ^40^. These positions are not conserved between HA types, or even within them. This suggests that despite many fixations at these positions and throughout the receptor-binding domain, the evolutionary constraints acting on the hemagglutinin protein have maintained receptor-binding pockets that are similar enough in structure and mechanism to be able to alter binding preference via similar mutations.

In this work, we draw on repeated patterns during independent evolutions of human influenza virus to chicken eggs to infer patterns underlying this adaptive landscape. Because of this, we are blind to strain-specific effects. For instance, if there is just a single egg-passaged strain available within a small clade of human virus, we will not infer mutations private to that egg-passaged strain to be adaptive, even though it is possible that they represent bona fide egg-adaptation that is specific to that rare background. This causes us to potentially underestimate the number of egg-adaptive pathways in areas of the human virus phylogeny where egg-passaged sequences are sparse. This is a compromise we make to be confident that the mutations we are analyzing have a true adaptive function.

Another limitation of this work is that there is variation in the number of times the different strains were passaged in eggs, and thus in the amount of evolutionary time they had to adapt. This could result in egg-passaged strains that are at an intermediate point in their adaptation (as hinted at in Figure 6). This complicates direct comparisons of individual strains, but again we draw power from comparing hundreds or thousands of strains. Given more passaging time, we would likely see a higher total number of strains with adaptive mutations, more mutational co-occurrence, and potentially a few more adaptive mutations with smaller (or more background-dependent) selection coefficients. However, this is unlikely to change the conclusions about egg-adaptation or its variance between subtypes that we discuss throughout this paper.

Here, we have looked to repeated adaptive evolution as a way of assessing the fitness landscape by using reiterated patterns of mutational co-occurrence to infer peakedness and epistatic constraints. Another common way to do this is by employing deep mutational scanning (DMS) to exhaustively determine the effect of every combination of mutations ^41–44^. This strategy has the benefit of directly measuring the effect of all pairwise mutation combinations, however it is limited both to phenotypes that are easily measurable in large-scale laboratory experiments and to a relatively small number of interacting sites. For instance, Wu et al ^45^ work out epistatic interactions between 11 residues in the HA receptor-binding site by quantifying the fitness effect of all pairwise combinations of amino acids, but extending this experiment to all of HA1 would be intractable (>19 million variants). With the goal of pushing our understanding of viral evolution past the effects of individual mutations on specific strains toward a more comprehensive view of the adaptive seascape the virus is evolving on, we view observation of natural evolutionary replicates as a complementary approach to exhaustive mutagenesis coupled with experimental phenotype measurement, with each method offering different advantages and insights.

## Acknowledgements

We gratefully acknowledge the authors who generated and submitted the viral sequences deposited in the NCBI GenBank database and the GISAID Initiative, on which the analyses in this manuscript are based. We have included an acknowledgements table for the sequences at https://github.com/blab/egg-adaptation/nextstrain_builds/gisaid_acknowledgements/gisaid_ackn owledgements.tsv.

## Methods

All code associated with the analyses in this manuscript is hosted at: https://github.com/blab/egg-adaptation.

### Sequence data and phylogenetic tree enriched in egg-passaged strains

Sequence data for all egg-passaged seasonal influenza strains was downloaded from GISAID separately for each segment (PB1, PB2, PA, HA, NP, NA, MP, NS) of seasonal influenza virus (H3N2, H1N1pdm, Vic and Yam) separately. We then filtered this data to include only one egg-passaged sequence per strain. Some strains that were passaged in eggs have been sequenced multiple times after different numbers of passages (and/or by different labs), and we chose to include the strain that was passaged the most number of times. We downloaded these sequences on March 31, 2025 and took only sequences between then and 30 years prior to this date. We then downloaded non-egg-passaged sequences for each segment and virus and subsampled them to ∼10000 sequences sampled as evenly as possible over this 30 year time window and over geographic regions. Within these sequences, we include all available non-egg-passaged sequences of each of the egg-passaged strains (referred to as “pairs” in Figure 1E). We combined the egg-passaged strains and the contextual non-egg-passaged strains together to form the dataset which we used to build the phylogeny with a Nextstrain pipeline. All sequences were aligned to a reference strain (H3N2 = A/Aichi/2/1968, H1N1pdm = A/California/07/2009, Vic = B/Russia/1969, Yam = B/Yamagata/16/1988) using Nextclade3. Trees were built using iqtree. During tree building, nucleotide sites that commonly mutate during egg-passaging were masked with an N so that they did not influence the topology. The masked sites (relative to the reference coordinates) were: H3N2 HA 556, 633, 634, 658, 750, and 751, H1N1pdm HA 631, 642, 735, 736, and 739, Vic HA 665, 666, 670, and 671, Yam HA 645, 646, 647, 651, and 652. All sites were then unmasked and TreeTime was used to estimate dates and sequences of internal nodes and assign mutations to them. Subsequent to exporting trees into auspice JSONs, we broke clusters of egg-only sequences into polytomies. These egg-enriched influenza trees can be viewed here https://nextstrain.org/groups/blab/h3n2/ha/egg, and the subtype and segment in the URL can be changed to view all 32 subtype and segment combinations. Acknowledgements for the labs that generated the sequences used in this study can be found in tabular format at https://github.com/blab/egg-adaptation/nextstrain_builds/gisaid_acknowledgements/gisaid_ackn owledgements.tsv.

### Egg-passaging mutation and adaptive substitution inference

All mutations that occurred during egg-passaging are inferred to be those occurring on terminal branches leading to egg-passaged strains. For each influenza virus and each segment, we store a list of all egg-passaging mutations that occurred for all egg-passaging strains inferred from the phylogeny in the folder ‘egg-mutation-analysis/egg-muts-by-strain/’. We also store a summary of the total number of egg-passaged strains, the frequency with an egg-passaged mutation, frequencies of all specific egg-passaged substitutions and of mutations to all residues, as well as a list of mutations that exceed 1%, 2.5%, and 5% of egg-passaged strains in ‘egg-mutation-analysis/egg-mut-counts/’.

We then infer probable adaptive substitutions for each segment of each subtype as those that occur independently in at least 5 different strains and where a) at least 2.5% of egg-passaged strains have an egg-passaging mutation at that residue, or b) at least 7.5% of egg-passaged strains within a 3-year window have that exact substitution. Adaptive mutations are calculated in ‘egg-mutation-analysis/get_adaptive_muts.ipynb’ and stored in JSON files in ‘egg-mutation-analysis/egg-adaptive-muts’.

### Protein structures

For visual representation of the location of egg-adaptive mutations on the hemagglutinin protein, we used solved structures of HA deposited on PDB with the accessions 2YP4 (H3N2) ^46^, 3UBE (H1N1pdm) ^47^, 4FQM (Vic) ^48^, and 4M40 (Yam) ^49^. For the nucleoprotein structures, we used 7NT8 (H3N2) ^50^ and 3ZDP (H1N1) ^51^.

### Time window analysis

The changing prevalence of egg-adaptive substitutions over time was assessed by apportioning the egg-passaged sequences into overlapping 3 year windows. Within each 3 year window, the proportion of strains with an adaptive amino acid substitution was determined for each residue position in HA1 with adaptive mutations. For each residue position with more than one adaptive amino acid substitution, the proportion of egg-passaged strains with a mutation to the different amino acids was calculated for each year. This analysis is done in ‘figure_panels/Figure3-timeWindows.ipynb’.

### Background genotype correlations with egg-adaptive mutations

For each temporally-restricted egg-adaptive mutation, we used the tree to identify HA1 mutations happening along trunk that precede the egg-adaptive mutation’s appearance, or follow the mutation’s disappearance. We then determined which of these trunk mutations correlate perfectly or nearly perfectly with strains containing the egg-adaptive mutation. If multiple trunk mutations had a good correlation with the egg-adaptive mutation, we presume mutations at the same residue are most likely to be responsible for the temporal restriction, and mutations at structurally proximal residues are the next most likely.

### Mutational co-occurrence enrichment score

For each influenza subtype, a mutational co-occurrence score was determined for every pair of adaptive mutations that overlap temporally. We define mutation A and mutation B as co-occurring temporally if there are at least 5 of each mutation and 25 A plus B mutations within the intersection of the date range over which A and B are observed. Mutation pairs that do not meet this criteria are considered non-overlapping and are colored grey.

For pairs of mutations that do overlap, we calculate the individual frequencies of mutation A and mutation B as well as the paired frequency of A and B in the time window of overlap. We then compute an enrichment score using the formula 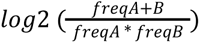. Pairs of mutations that never co-occur result in a mathematically undefined value, but represent a severe depletion of co-occurrence. We color these pairs dark blue.

For this analysis, we separate the different substitutions at a given position if the amino acid substitutions change over time (see time window analysis). For instance, H3N2 186S, 186V, and 186N are considered different adaptive mutations for this analysis because they occur during different periods of time, but 156Q and 156R are grouped together because they occur during the same period of time. The code for this analysis is in ‘figure_panels/Figure5-betweenMutEpistasis.ipynb’.

### Sets of egg-passaged strains from identical HA1 sequences

For each egg-passaged strain, the HA1 sequence before egg-passaging is inferred as the sequence at the first internal node in the phylogeny. This is done by applying every mutation along the path between the root and this node to the root sequence. For each strain, the inferred sequence before egg-passaging is stored in a subtype-specific file like ‘egg-mutation-analysis/before-egg-seqs/h3n2_HA1_seqs_before_egg.json’. All egg-passaged sequences with identical HA1 sequences prior to egg-passaging are determined and grouped into sets of “replicates”, which are stored in ‘egg-mutation-analysis/before-egg-seqs/h3n2_replicate_sets.json’. For each set, a “haplotype” is assigned to each constituent strain within the set, where haplotype refers to which egg-adaptive sites (residues) are mutated in that strain. The number of strains with each haplotype are counted to produce the plots in Figure 6A and Supplemental Figures 4-7, representing egg-adaptation of every replicate within a set.

### Avian H1 and H3 phylogenies

All alphainfluenzavirus (NCBI:txid197911) sequences from an avian host were downloaded from NCBI Genbank in FASTA format on June 4, 2024 (n=271,324). HA type was parsed from the metadata, where available, using the custom script ‘parse_ncbi_fasta.py’, which outputs one FASTA data file per HA type. Using the H1 and H3 files as input, we then ran a Nextstrain pipeline to filter, subsample, and align sequences, and to reconstruct a time-resolved phylogeny and infer mutations across the tree. Specifically, we used ‘augur filter’ to select only hemagglutinin sequences, and to subsample a maximum of 50 samples per year between 1950 and the present. We used ‘augur align’ to align all H1 sequences to A/California/07/2009 (accession: CY121680) and all H3 sequences to A/Aichi/2/1968 (accession:CY121117). These are the same reference sequences we used to generate the human and egg-passaged H1N1pdm and H3N2 trees used throughout this work, ensuring that the coordinate systems are consistent. We then build trees using IQ-TREE and use Treetime to estimate molecular clock branch lengths, rooting on an outgroup sequence. We root H1 on the avian H3 A/mallard/Wisconsin/418/1976 HA sequence (accession: CY180420) and H3 on the avian H1 A/widgeon/Wisconsin/505/1979 HA sequence (accession: CY179395). We prune the outgroup sequence and then reconstruct mutations along branches using Treetime. The resulting trees of 901 avian H1NX (https://nextstrain.org/groups/blab/avian/H1) and 1284 avian H3NX strains (https://nextstrain.org/groups/blab/avian/H3) are saved in Auspice format and are viewable at the links provided.

### Human and avian genotype analysis

The predominant amino acids observed amongst human H3N2 and H1N1pdm strains at each egg-adaptive site were determined from all non-egg-passaged strains in the phylogenies used throughout this paper. The avian H3NX and H1NX trees described above were used to determine predominant avian genotypes at these positions. All amino acids observed in 10% or more of the strains were recorded.

