## Supplemental Figures for "Seasonal influenza viruses show distinct adaptive dynamics during growth in chicken eggs"

### Supplements

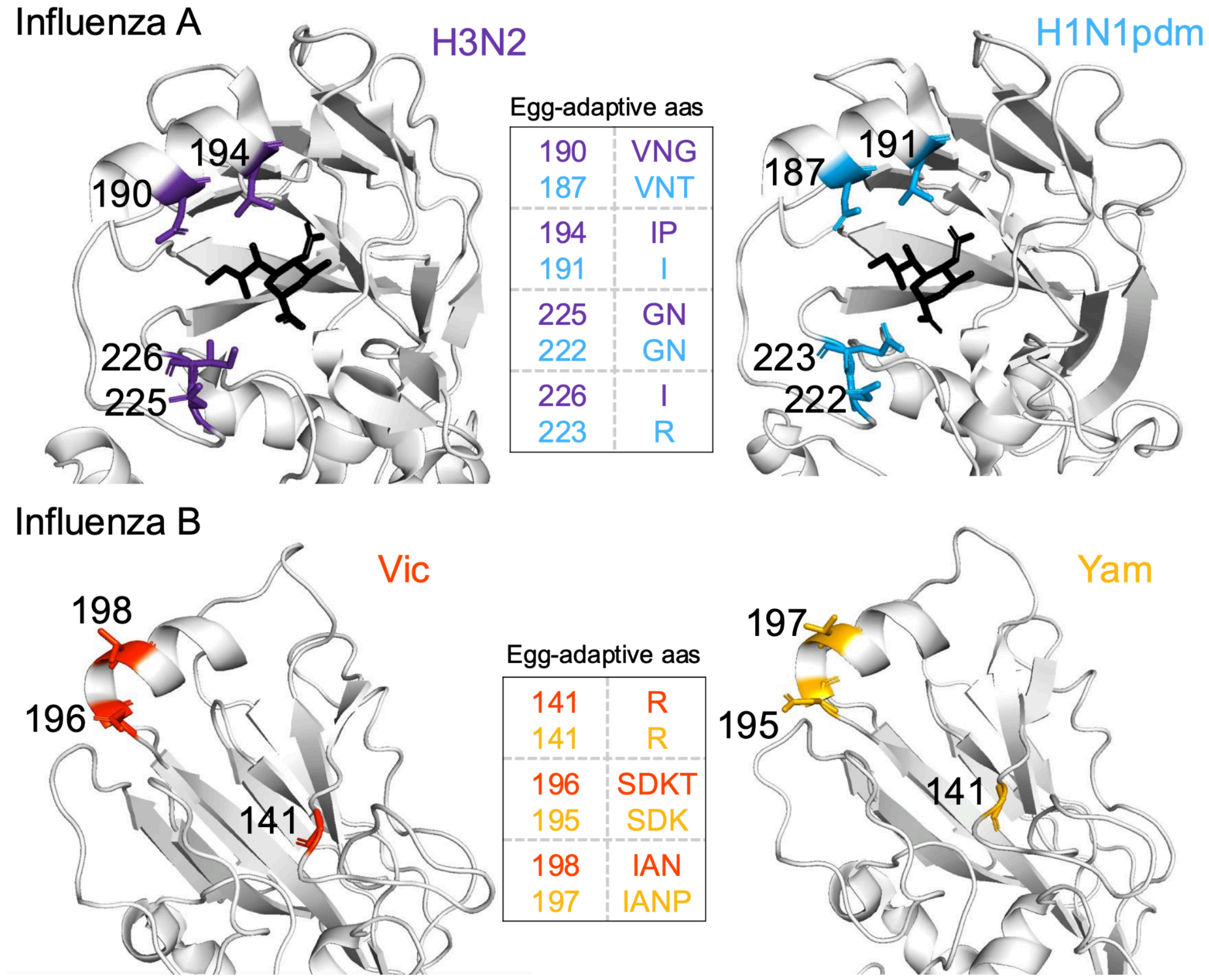


**Supplemental Figure 1. Structurally homologous locations of egg-adaptive mutations in HA1.** Residues that get egg-adaptive mutations in both influenza A viruses (top row) or both influenza B viruses (bottom row) are highlighted on the structure of HA. Egg-adaptive amino acids observed at these homologous sites in both viruses are listed in the table. Influenza A viruses are shown with the terminal moeity of an α-2,6 sialic acid analog (black) for context. The PDB for structures are H3N2 (2YP4), H1N1pdm (3UBE), Vic (4FQM), Yam (4M40).

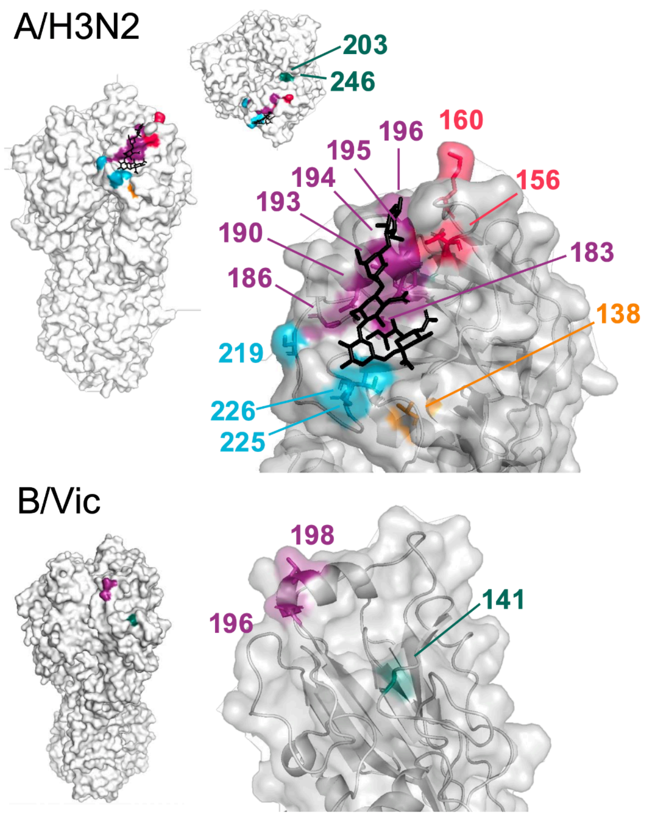


**Supplemental Figure 2. Locations of all HA1 egg-adaptive mutations.** The position of all egg-adaptive mutations in HA are highlighted on the protein structure for each influenza subtype. The PDB for structures are H3N2 (2YP4), H1N1pdm (3UBE), Vic (4FQM), Yam (4M40).

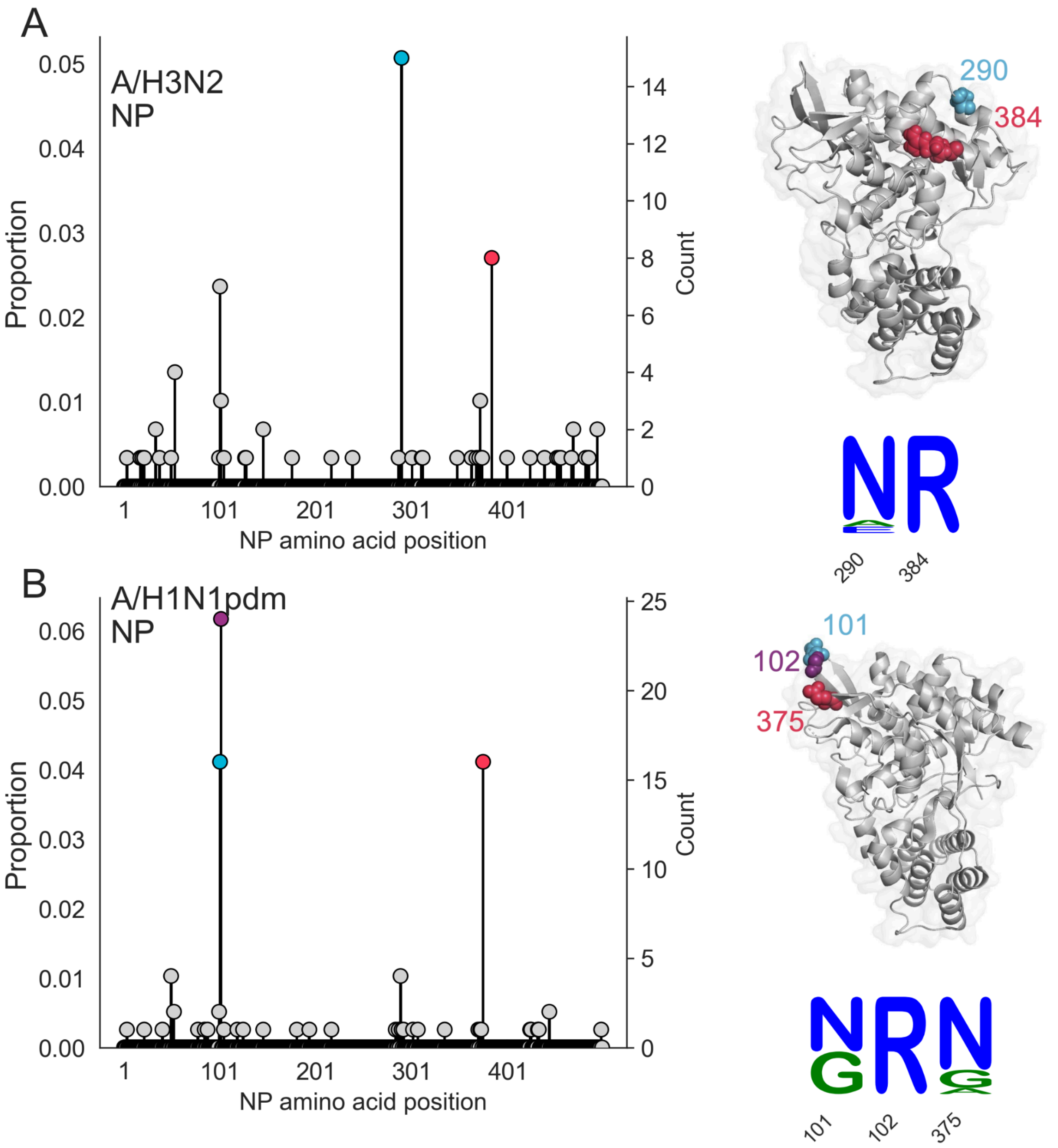


**Supplemental Figure 3. Egg-adaptive mutations in nucleoprotein of influenza A viruses.** Proportion of egg-passaged strains with a mutation at each residue in the NP protein for A) H3N2, and B) H1N1pdm. Inferred egg-adaptive mutations are colored and these same colors are used to indicate the position of each mutation on the structure of NP. The H3N2 structure is from a 1968 strain (PDB 7nt8), and the H1N1 structure is from the 1933 WSN H1N1 strain (PDB 3zdp). Logo plots show the amino acid substitutions at each egg-adaptive site in NP, in proportions.

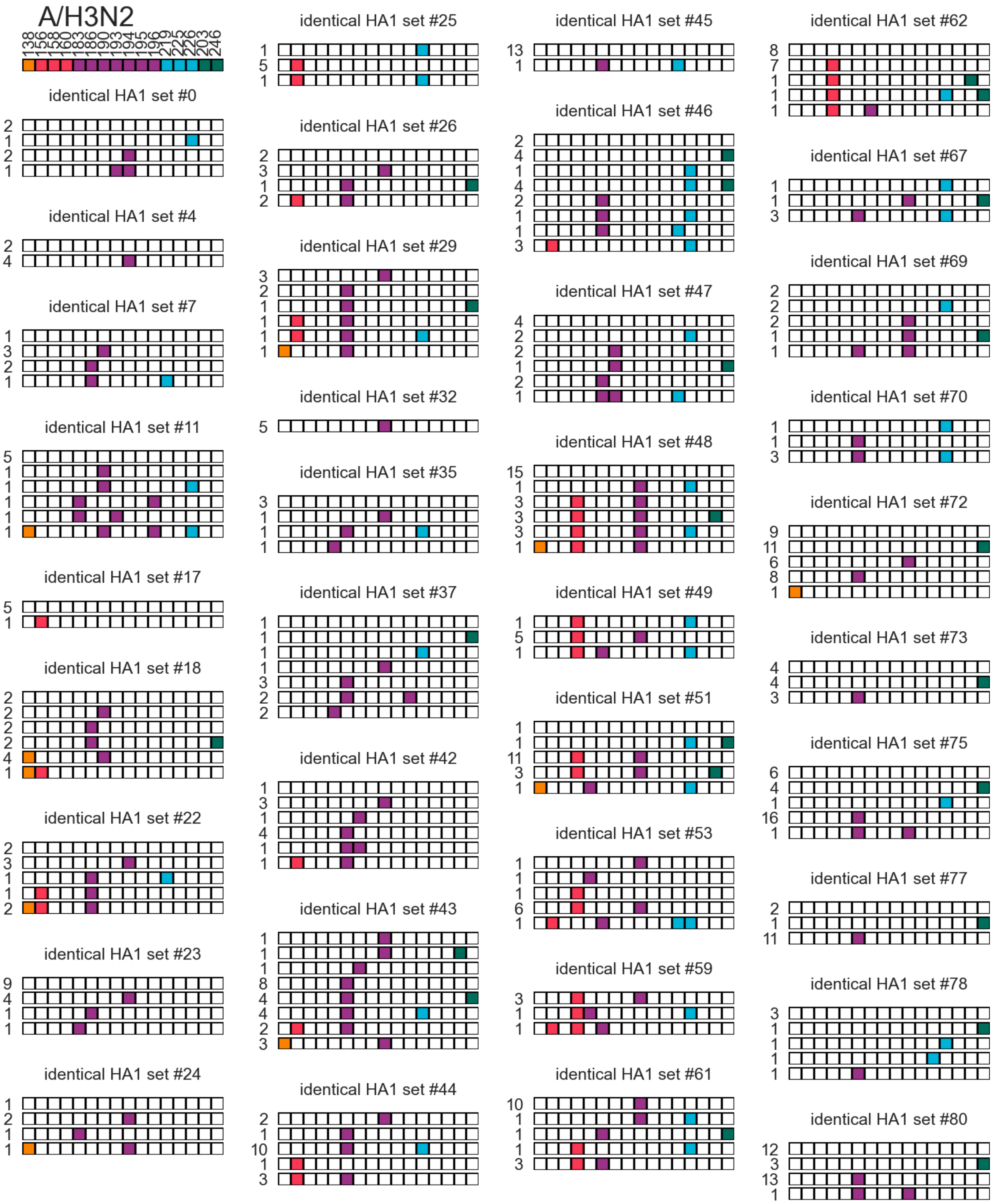


**Supplemental Figure 4. All sets of A/H3N2 egg-passaged strains derived from identical HA1 sequences.**

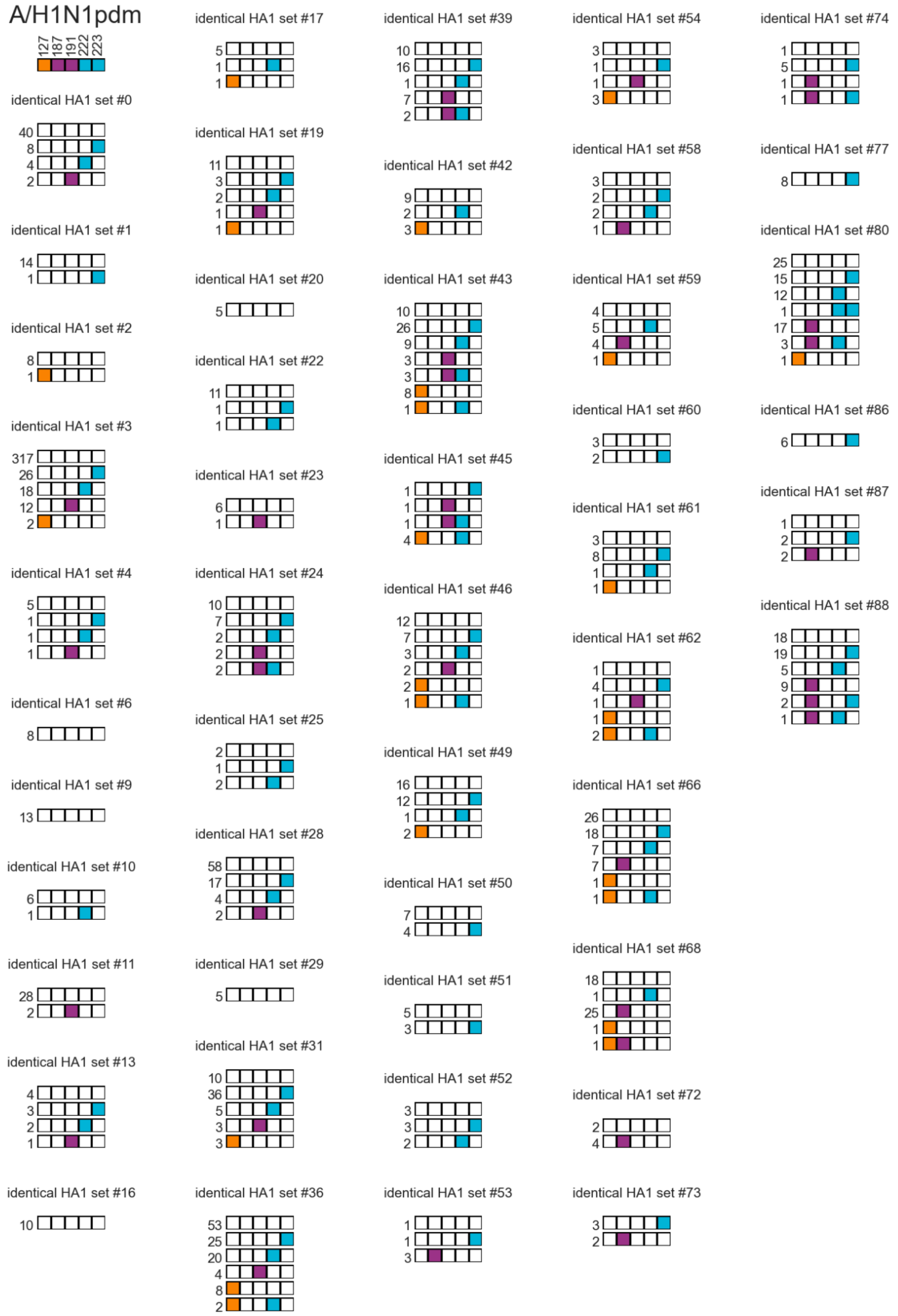


**Supplemental Figure 5. All sets of A/H1N1pdm egg-passaged strains derived from identical HA1 sequences.**

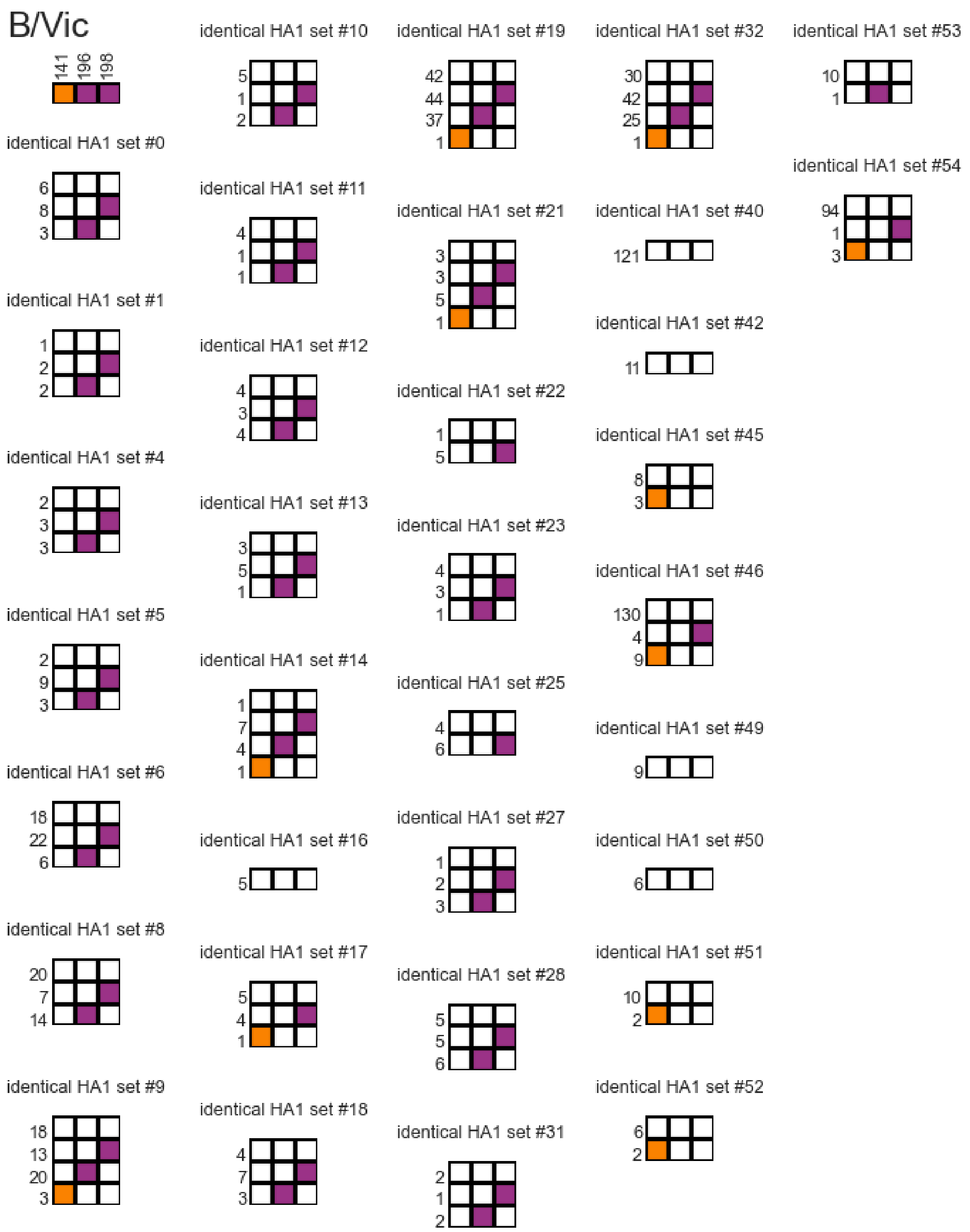


**Supplemental Figure 6. All sets of B/Vic egg-passaged strains derived from identical HA1 sequences.**

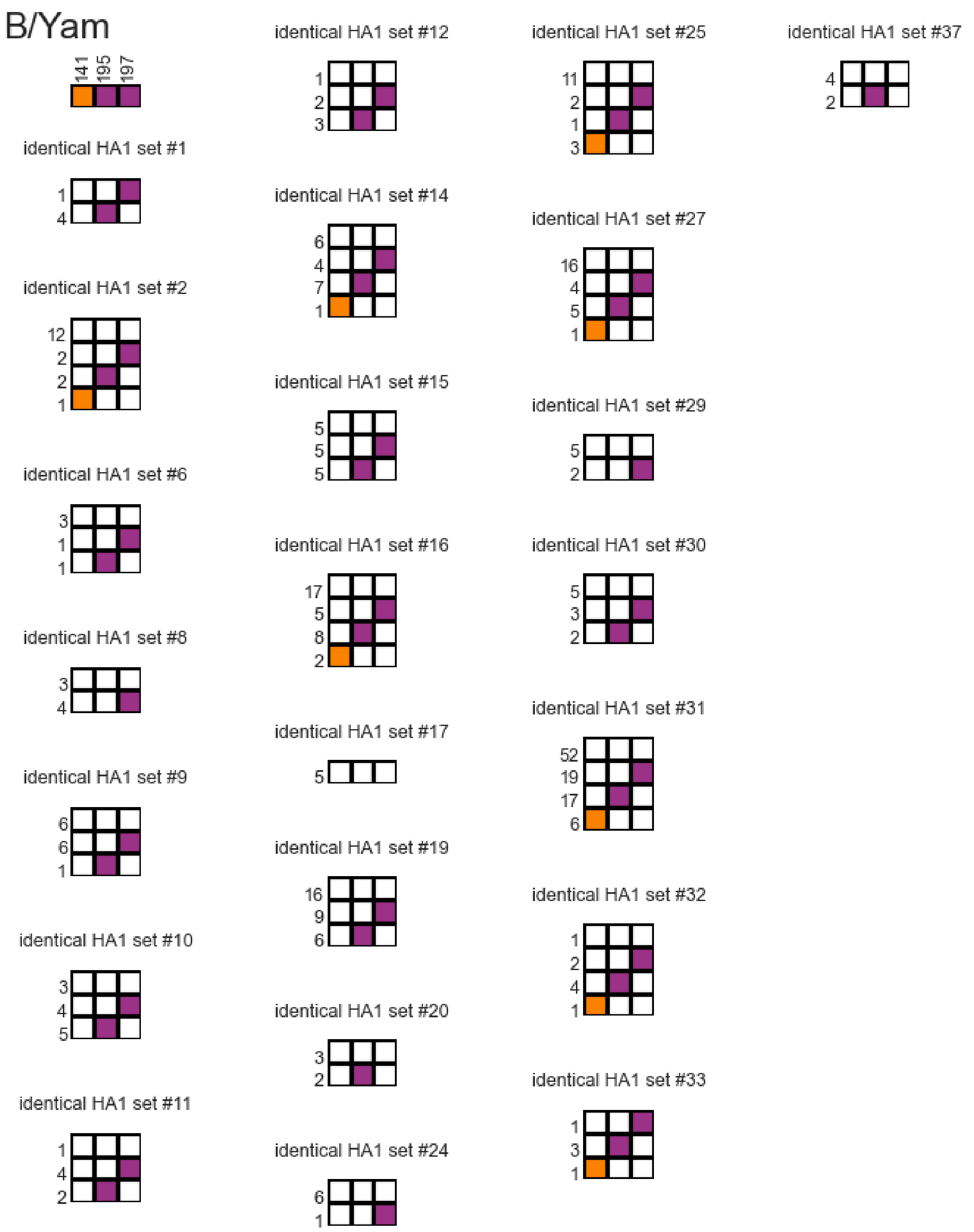


**Supplemental Figure 7. All sets of B/Yam egg-passaged strains derived from identical HA1 sequences.**

## A/H3N2

| | 138 | 156 | 160 | 183 | 186 | 190 | 193 | 194 | 195 | 196 | 203 | 219 | 225 | 226 | 246 |
|---|---|---|---|---|---|---|---|---|---|---|---|---|---|---|---|
| Human | A | H,Q,S | K,T,I | H | S,G,D | D,N | S,F | L | Y,F | V,A | T | S | G,D,N | V,I | N |
| Egg | S | Q,R | K,I | L | V,N,S | N,G,V | R | P,I | Y | T | I | Y,F | G,N | I | K,H,S,T |
| Avian | A | K | A,T | H | S | E | N,S | L | Y | V | T | S | G | Q | N |

#### A/H1N1pdm

| | 127 | 187 | 191 | 222 | 223 |
|---|---|---|---|---|---|
| Human | D | D | L | D | Q |
| Egg | E | V,N,T | I | G,N | R |
| Avian | E | E | L | G | Q |

**Supplemental Figure 8. Egg-adaptive mutations make human viruses more avian-like at some, but not all, adaptive sites.** Amino acids observed at each egg-adaptive site in the human HA, egg-adapted human HA, and avian HA. For the human and avian HA, all amino acids observed in 20% or more of the sequences are listed.

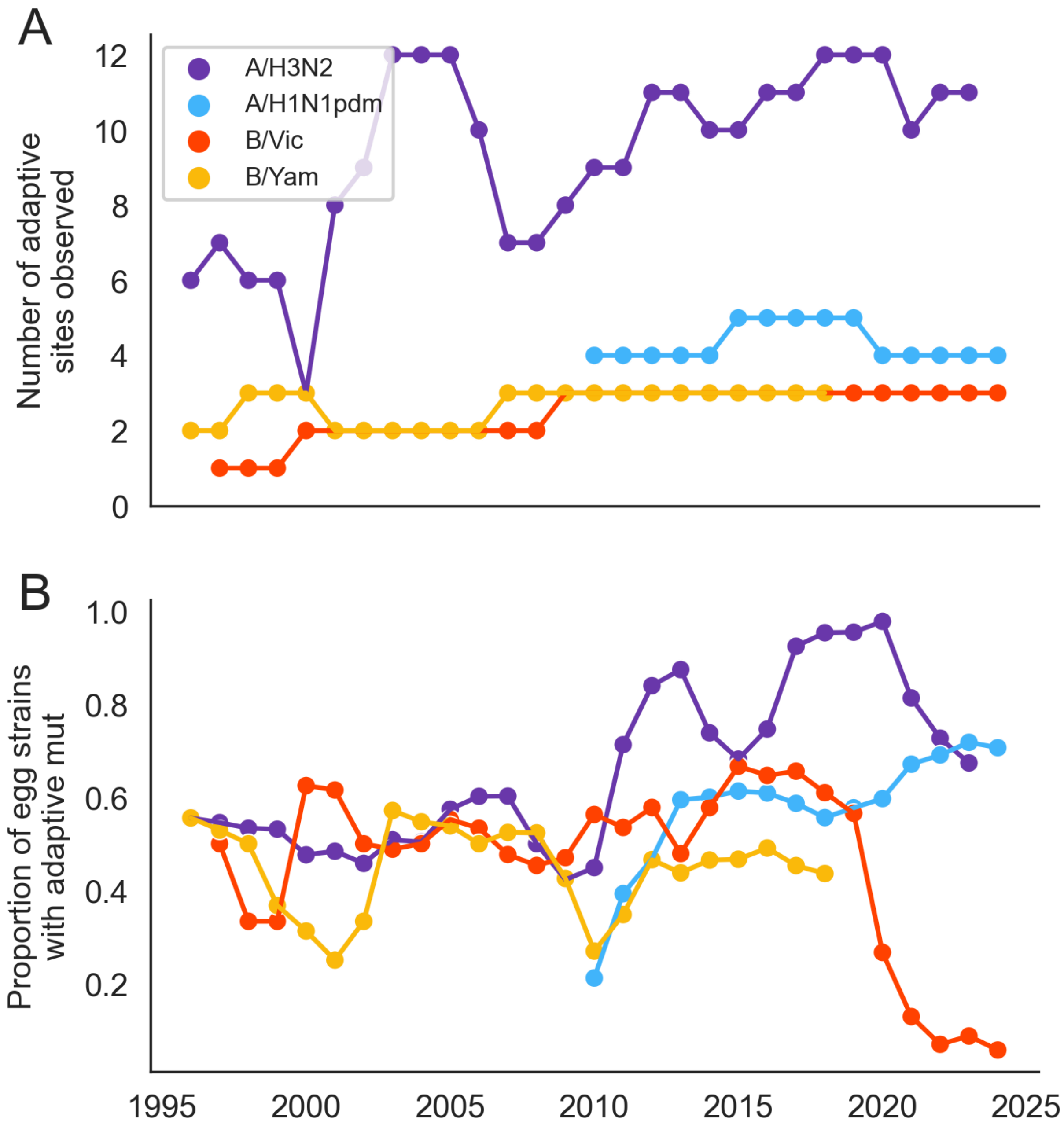


**Supplemental Figure 9. Total number of egg-adaptive mutations differs more over time for H3N2 than for H1N1pdm or influenza B viruses.** A) The number of egg-adaptive sites (residues) in HA where a mutation is observed in at least 2 strains or at least 10% of strains during a 3-year window. B) The proportion of egg-passaged strains within a 3-year window that have an adaptive mutation in HA. Both panels are calculated in sliding 3-year windows, plotted at the midpoint year.
